# Generating dynamic carbon-dioxide from the respiratory-volume time series: A feasibility study using neural networks

**DOI:** 10.1101/2022.07.11.499585

**Authors:** V. Agrawal, Xiaole Z. Zhong, J. J. Chen

**Affiliations:** Rotman Research Institute, Baycrest Centre for Geriatric Care, Toronto, Canada; Department of Medical Biophysics, University of Toronto, Canada

**Keywords:** Carbon dioxide (CO_2_), end-tidal CO_2_, respiration, respiratory volume, respiratory volume variability (RVT), respiratory response function (RRF), resting-state fMRI, cerebrovascular reactivity, physiological recordings, deep learning, convolutional neural network (CNN), fully convoluted neural network (FCN)

## Abstract

In the context of fMRI, carbon dioxide (CO_2_) is a well-known vasodilator that has been widely used to monitor and interrogate vascular physiology. Moreover, spontaneous fluctuations in end-tidal carbon dioxide (PETCO_2_) reflects changes in arterial CO_2_ and has been demonstrated as the largest physiological noise source in the low-frequency range of the resting-state fMRI (rs-fMRI) signal. Increasing appreciation for the role of CO_2_ in fMRI has given rise to methods that use it for physiological denoising or estimating cerebrovascular reactivity. However, the majority of rs-fMRI studies do not involve CO_2_ recordings, and most often only heart rate and respiration are recorded. While the intrinsic link between these latter metrics and CO_2_ led to suggested possible analytical models, they have not been widely applied. In this proof-of-concept study, we propose a deep learning approach to reconstruct CO_2_ and PETCO_2_ data from respiration waveforms in the resting state. We demonstrate that the one-to-one mapping between respiration and CO_2_ recordings can be well predicted using fully convolutional networks (FCNs), achieving a Pearson correlation coefficient (r) of 0.946 ± 0.056 with the ground truth CO_2_. Moreover, dynamic PETCO_2_ can be successfully derived from the predicted CO_2_, achieving r of 0.512 ± 0.269 with the ground truth. Importantly, the FCN-based methods outperform previously proposed analytical methods. In addition, we provide guidelines for quality assurance of respiration recordings for the purposes of CO_2_ prediction. Our results demonstrate that dynamic CO_2_ can be obtained from respiration-volume using neural networks, complementing the still few reports in deep-learning of physiological fMRI signals, and paving the way for further research in deep-learning based bio-signal processing.

## Introduction

Carbon dioxide (CO_2_) is a potent vasodilator used that has been shown to rely mainly on the nitric oxide pathway to increase arterial diameter (Iadecola, 2017; Najarian et al., 2000; Peebles et al., 2008; Pelligrino et al., 1999). Blood-vessel diameter is highly sensitive to the surrounding CO_2_ concentration, with increasing CO_2_ partial pressures leading to linear increases in both vessel diameter and flow (Hülsmann and Dubelaar, 1988; Komori et al., 2007). In Komori et al. for example, this increase was shown to be 21.6% for arteriolar diameter and 34.5% flow velocity for a 50% change in CO_2_ partial pressure in rabbit arterioles (Komori et al., 2007). The partial pressure of carbon dioxide (PCO_2_) is the measure of CO_2_ within arterial or venous blood. It often serves as a marker of sufficient alveolar ventilation within the lungs. Under normal physiologic conditions, the value of PCO_2_ ranges between 35 to 45 mmHg, or 4.7 to 6.0 kPa. Typically the measurement of PCO_2_ is performed via arterial blood gas, but the end-tidal pressure of CO_2_ (PETCO_2_) is related to intravascular PCO_2_ through a linear relationship under steady-state conditions (Peebles et al., 2007, 2008), allowing arterial PCO_2_ to be estimated from PETCO_2_.

Dynamic CO_2_ recordings have multiple utilities and implications. In the past decades, the CO_2_-driven functional magnetic resonance imaging (fMRI) response has been the preeminent method for mapping cerebrovascular reactivity (Blockley et al., 2017; Chen, 2018; Chen and Gauthier, 2021). Wise et al. first reported the contribution of spontaneous fluctuations in arterial PCO_2_ to the resting-state fMRI (Wise et al., 2004). Chang et al. followed up this work by demonstrating the potential relationship between PETCO_2_ and respiratory variability (RVT) (Chang and Glover, 2009). Using recordings of spontaneous PETCO_2_ variations, Golestani et al. determined the fMRI response function that links PETCO_2_ to the resting-state blood-oxygenation level dependent (BOLD) signal (Golestani et al., 2015), and also demonstrated PETCO_2_ as the primary source of physiological noise in resting-state BOLD. It has even been used to demonstrate the possible existence of neuronally-motivated vascular networks in the brain (Bright et al., 2020). Furthermore, Chan et al. found that PCO_2_ (not PETCO_2_) fluctuations also contribute significantly to resting-state BOLD signal variability (Chan et al., 2020, n.d.). While the mid-breath PCO_2_ does not reflect intravascular PCO_2_, PETCO_2_ does provide a quantitative estimate of arterial PCO_2_, and is more widely used in fMRI experiments for the purposes of denoising (Murphy et al., 2013) and CVR mapping (Pinto et al., 2020). The substantial influence of dynamic PETCO_2_ fluctuations on resting-state (Golestani and Chen, 2020) and dynamic functional connectivity has been demonstrated recently (Nikolaou et al., 2016). Dynamic CO_2_ can also allow vascular lag structures to be estimated, providing an important metric for assessing vascular health (Champagne et al., 2019). Given the unique variance explained by PCO2 and PETCO_2_, it is safe to say that dynamic CO_2_ is a useful thus desirable metric for those working with resting-state fMRI data.

Despite the increasing realization of the value of CO_2_ recordings, it is often impossible to obtain recordings of CO_2_ during an fMRI session. Most study sites are not equipped with an MRI-compatible capnometer that also facilitates continuous recording of PCO_2_. Moreover, the many thousands of legacy fMRI data sets (e.g. Human Connectome Project, UK Biobank) certainly do not include CO_2_ recordings. On the other hand, respiratory volume variations, which had previously been related to PETCO_2_ variations, are more readily available thanks to the incorporation of respiratory-volume belts in modern MRI systems. RVT was first introduced by Birn et al. as a noise source in fMRI that introduces unique signal variability (Birn et al., 2006). Today, while RVT measurements during fMRI sessions are increasingly common, they are still unavailable in large-scale studies and legacy data sets. As a possible solution, recent work by Salas et al. (Salas et al., 2020) demonstrated that the RVT time series can in principle be reconstructed from fMRI data using a convolutional neural network (CNN).

Chang et al. previously showed that PETCO_2_ can be related to RVT through convolution with a respiratory-response function (Birn et al., 2008). However, this relationship has been difficult to reproduce in resting-state conditions, as we will show with our data. In the resting state, not only is it impossible to derive quantitative CO_2_ values from respiratory volume, it is also difficult to obtain a deterministic relationship between dynamic patterns of respiratory volume and CO_2_ variation. Thus, in this study, we also use the principle of deep learning, but our focus is to bridge the gap between respiratory and CO_2_ recordings. Our aim is to demonstrate the feasibility of using deep learning to produce dynamic CO_2_ waveforms from the respiratory time series.

## Background on Neural Networks

In the majority of deep learning studies in neuroimaging applications, 2D inputs are used to produce 2D outputs (Zhu et al., 2019). Image-to-image translation is used for cross-modality conversion, denoising, super-resolution and reconstruction (Kaji and Kida, 2019). Our problem entails the estimation of a 1D signal from another 1D signal. Within this context, past research has used CNNs and recurrent neural networks (RNNs). Traditional convolutional neural networks (CNNs) consist of convolutional layers followed by fully connected layers (dense layers) at the end of the architecture (Rawat and Wang, 2017). As CNNs are the most successful type of deep-learning model for 2D image analysis, and physiological signals are 1D time-series data, some have converted 1D signals to 2D data to be fed into a CNN network, and have obtained good results (Zhang et al., 2020). The advantage of using 1D CNNs over 2D CNNs and RNNs is the significant reduction of the number of training parameters, which is helpful when the training data is limited. Applications of 1D CNNs typically include ECG classification and anomaly detection biomedical data classification (Kiranyaz et al., 2021). Salas et al. pioneered the use of 1D CNN for estimating physiological fluctuations in fMRI and is closely related to our domain. They segmented BOLD fMRI signals into fixed time-windows and fed them into a CNN, where the dense layer predicts a single point estimate of the respiration waveform at the center of the window. To predict the entire time series, all the mini time-windows have to be separately propagated through the network, increasing the complexity and computational cost of the approach. Moreover, due to the presence of the dense layer, traditional CNNs cannot be trained for variable length input-output regression tasks.

In this work, we have implemented a specific type of CNNs known as fully convolutional networks (FCNs) (Long et al., 2015). FCNs are nothing but a traditional CNN without fully connected layers. Fully convolutional layers in FCN permits the use of variable length input and also minimizes the computational cost of the dense connections. Here, we extend FCNs’ image-to-image translation use case in 1D domain, wherein, the encoder-decoder architecture exploits the latent space to streamline the translation. Previously, 1D U-net (a type of FCN which includes skip connections) has been implemented for reconstructing low-frequency respiratory volume signals from fMRI (Bayrak et al., 2020). Skip connections as used in the U-net could be implemented in this study, but as the study is more focused on establishing the proof of concept, such complications were avoided in our implementation of FCNs.

## Methods

### Data Acquisition

We recorded percent-CO2 (%CO2) fluctuations and respiratory bellows simultaneously in a group of 18 healthy adults (age 20-38 years) using the Biopac System (Biopac Inc., Goleta, CA, USA). The Biopac respiration belt was positioned below the ribcage, and detects respiratory depth by sensing abdominal circumference changes. %CO_2_ data were acquired through gas lines attached to masks affixed to subjects’ faces. The Biopac %CO_2_ module (CO2100C) is calibrated to measure %CO_2_ concentration in the range of 0 to 10%. In total, the available data set consisted of 136 resting-state recordings from different subjects, which were 10.8 minutes long on average (min = 7.2 min, max = 16.1 min). To the best of our knowledge, this is the largest data set of its kind in existence.

### Data Preprocessing

The preprocessing steps consist of (1) low-pass filtering both respiration and CO_2_ waveforms (f < 1 Hz) and (2) correcting the delay between %CO_2_ and respiration signal by cross-correlation. The low pass filter’s cutoff frequency was determined based on the respiratory rate of an individual (0.2 - 0.4 Hz). The delay between %CO_2_ and respiration waveforms was corrected by shifting the %CO_2_ time course by the time lag yielding the maximum negative cross-correlations between it and the respiration waveform. We found that across all cases, to achieve this, the %CO2 time course had to be shifted to the left (backwards in time) by an average of 8.5 s (with a standard deviation of 1.5 s).

After the delay correction process, we rejected data that yielded absolute Pearson correlations of less than 0.4. Recordings were also rejected if their length was less than 3 minutes, too short to allow adequate training. More details on the correlation and data-length threshold are given in the quality assurance section. The respiration belt data was in arbitrary units, hence it was normalized by subtracting the temporal mean and dividing the result by standard deviation. The same procedure was applied to the %CO_2_ waveforms. Further details about the normalization are provided in the next subsection. Both the waveforms were then resampled to 10 Hz and exported in CSV format to be later imported during the training phase of the neural network.

To obtain PETCO_2_ from the normalized %CO_2_ recordings, the peak detection algorithm ensures the minimum distance between the two peaks is twice the sampling frequency. In other words, we assumed the time between two exhales is at least 2 seconds, which is consistent with our recorded respiratory intervals (3-5 s per breath). Moreover, the lower limit of the amplitude of the peak was set to be 0.3. It rejected any peaks which were either negative or too close to zero, thus, removing false positives.

#### Data Normalization

As previously mentioned, both %CO_2_ and respiration-belt data were demeaned (zero mean) and normalized to unit standard deviation (such that SD = 1). The respiration data is fluctuations in voltage transduced from expansions and contractions of the belt. As such, it varies with slight variations in belt tightness and positioning, and needs to be normalized across subjects to achieve inter-subject consistency. In part due to the need of using normalized respiration as the independent variable, this latter would encode no quantitative %CO_2_ information. That is, there could be a many-to-one relationship between normalized respiration and unnormalized CO_2_. To mitigate this issue, we demeaned and normalized the %CO_2_ time series in the same manner. In this manuscript, all the further mentions of CO_2_ denote normalized %CO_2_, unless stated otherwise.

#### Quality Assurance

A critical part of successful application of machine learning is quality assurance (QA) of the training and testing data.It is more probable to find noise in respiration data, wherein artifacts such as subject movement and talking can easily confound respiratory-belt recordings. Moreover, if the participant does not consistently breathe from the abdomen, the respiration belt data may not correspond well with the CO_2_ data. During the data-collection phase, useful precautions include ensuring that the respiration belt and CO_2_ gas lines are properly connected. Such precautions not only reduce the unwanted waveforms but also increase the feasibility of machine-learning approaches. To discard the undesirable recordings, we have evaluated our data based on the factors given below.

1. Length of the recording
2. Pearson correlation coefficient between CO_2_ and respiration after delay correction process
3. Low-frequency noise present in the waveforms

Nonetheless, it is informative to use data containing some level of noise and artifact for the purposes of representativeness. Therefore, the threshold used in the rejection process is generously selected.

##### Length of the recording

In general, for our approach, longer data sets are more desirable. It was observed that all the recordings were either less than 3 minutes or more than 6 minutes in length, drawing a clear distinction between test recordings and usable recordings. Thus, the lower limit for the time length was set to 3 minutes. **Figure 1** shows the histogram plot of all the recordings after the time-length thresholding.

**Figure 1.**
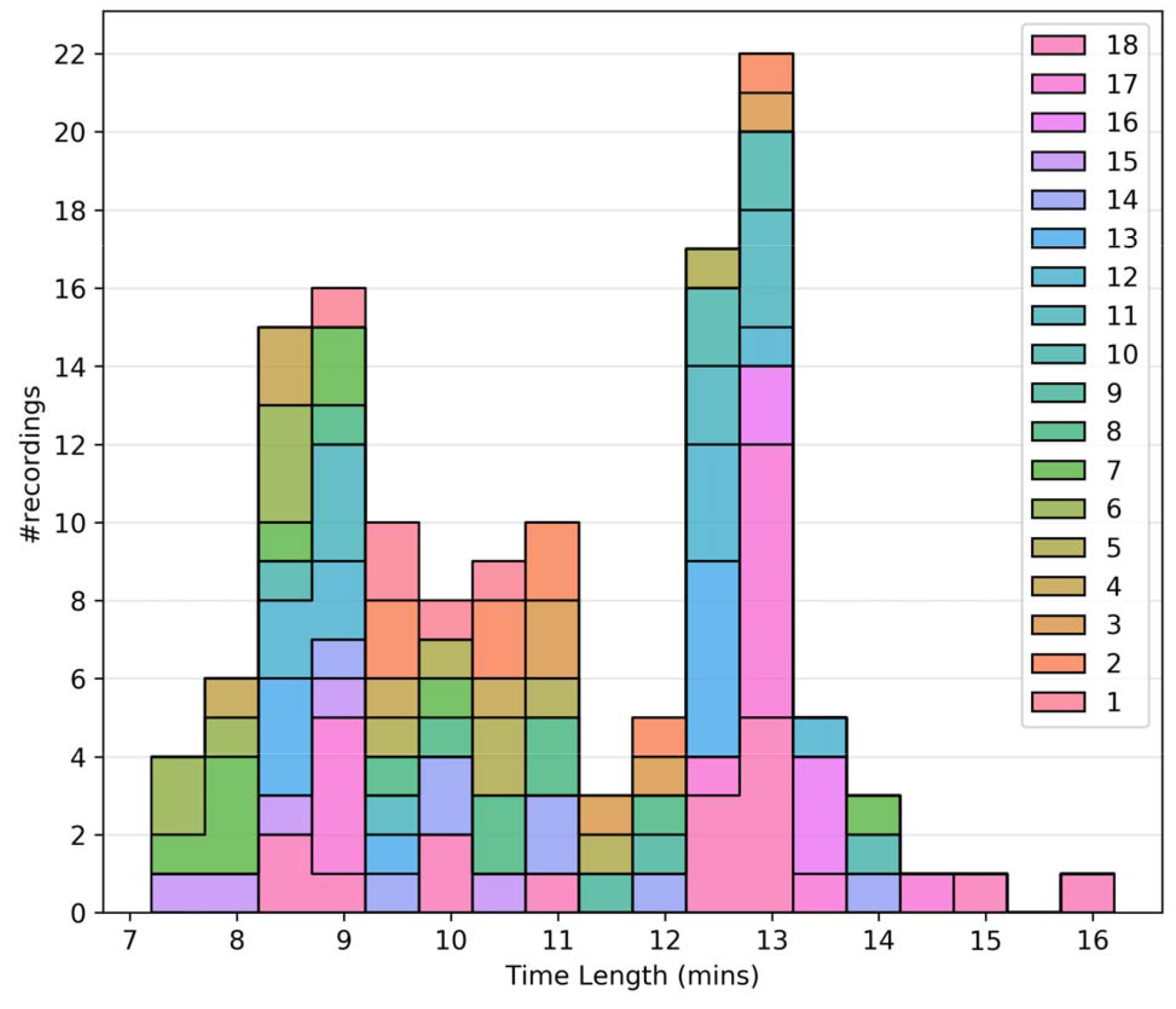
Quality assurance metrics: histogram plot of the time length of recordings after time length thresholding. Different colours are used to separate the subjects.

##### Pearson correlation coefficient

As previously mentioned, Pearson’s correlation (r) between the respiratory belt and CO_2_ time courses is used for initial QA purposes. The threshold for the absolute value of correlation between CO_2_ and respiration is -0.4, as respiratory volume and CO_2_ are expected to be negatively associated. This limit was empirically determined through manual review of the recordings. **Figure 2** shows that even though the threshold was -0.4, there were no recordings with r between -0.4 to -0.5, only one recording with r = -0.5 and most of the recordings had an r value of <-0.6.

**Figure 2.**
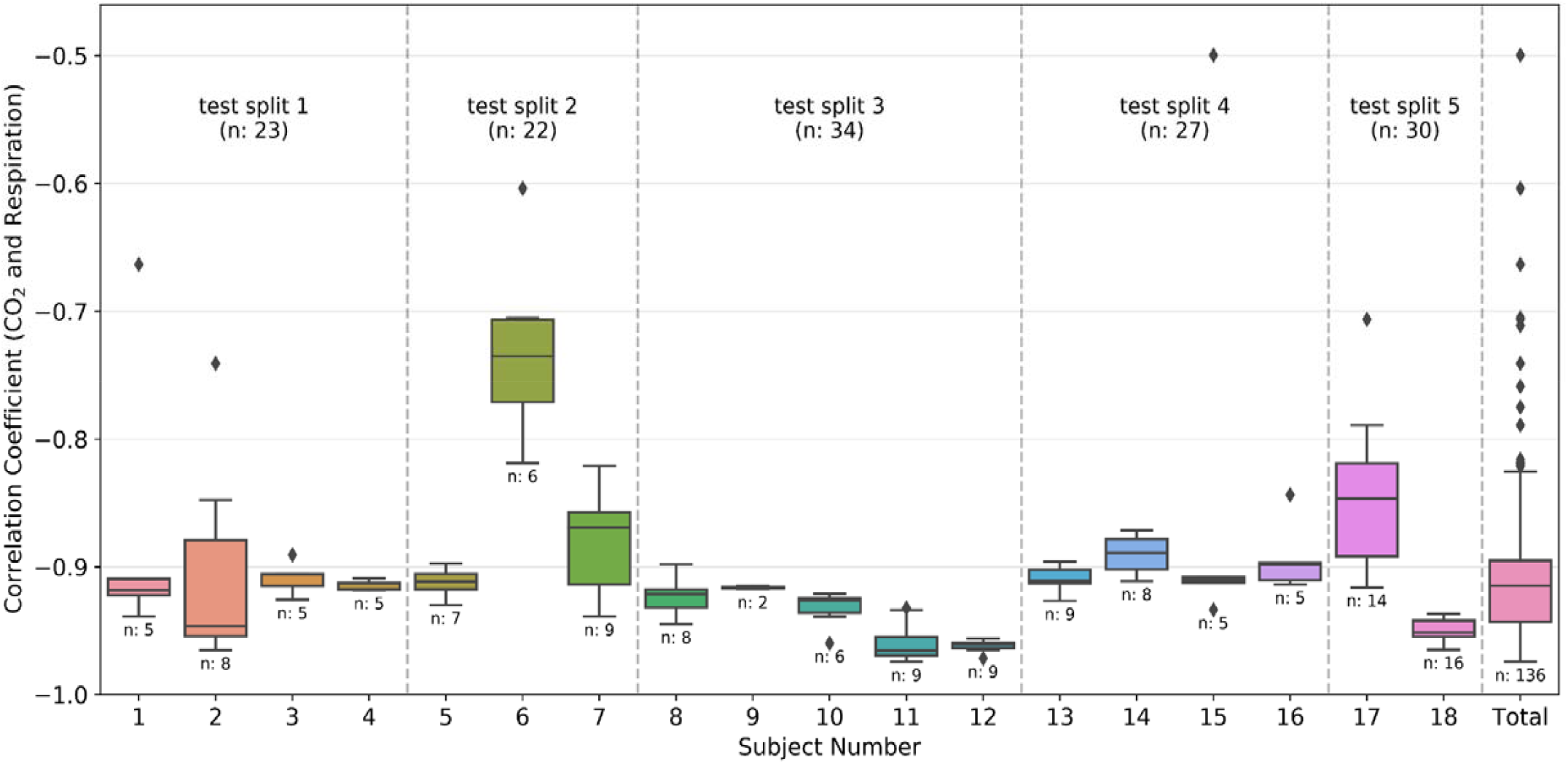
Quality assurance metrics: box plots of the correlation coefficient between CO2 and respiration waveforms from each individual subject and the total data after preprocessing. The number of recordings available for each subject is also given below the box plot. The divisions created by the dashed line show the groups made during the k-fold split of the dataset. The group number is the same as the test split number, and the total number of recordings in the group is also provided in the plot. The colour coding is the same as figure 1.

##### Low-frequency noise

Within the 0.1-0.5 Hz frequency band, noise in the respiratory and CO_2_ waveforms can impair our ability to relate the two waveforms, even if the recording-duration and correlation-coefficient thresholds are met. Such noise most likely originates from faulty attachment of the respiratory belt and from drifts in the recording modules. As it could potentially overlap with breathing frequency, it cannot be separated from the signal by using filters. However, this type of noise can be identified through a mismatch in the low-frequency portion (< 0. 2Hz) of the power spectra of CO_2_ and respiration, as shown in **Figure 3**. This type of noise is also reflected in the signal time series as periodic decreases or increases in the amplitude of signal. An example of this behaviour is shown in **Figure 3**, whereby the amplitude of the valleys in the respiratory data segmented by the red box significantly decreases or increases for multiple consecutive breaths, whereas this pattern remains unreflected in the CO_2_ waveform. This is contrasted with a sample usable data, shown in **Figure 4**.

**Figure 3.**
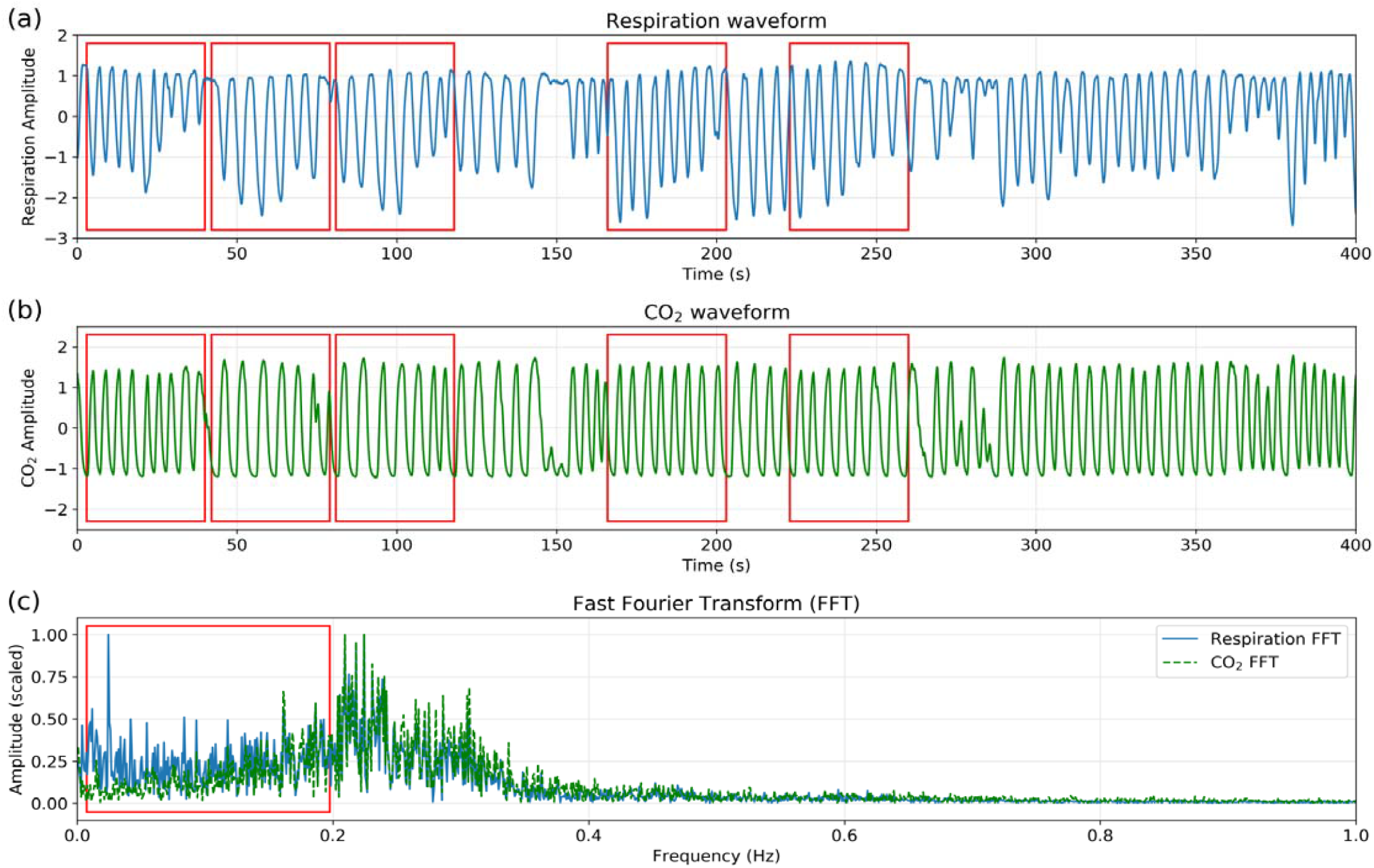
Quality assurance: sample data with low-frequency noise. (a) and (b) show a 400 seconds segment of the normalized respiration and CO_2_ recordings, respectively. In (c), the power spectra are shown. Red boxes superimposed on the plot are used to show the low frequency noise present in the data.

**Figure 4.**
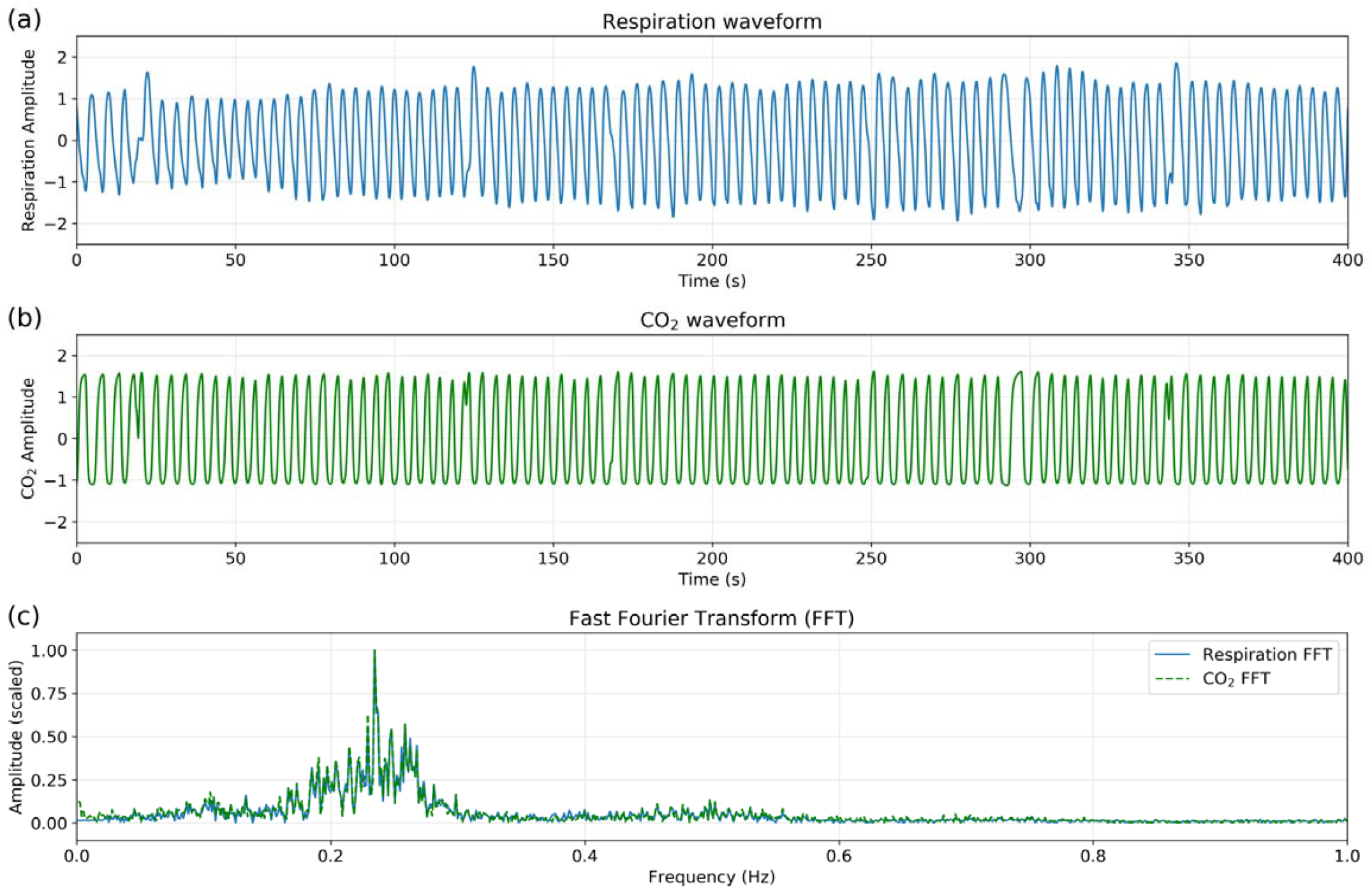
Quality assurance: sample clean data. Segment of a recording from our dataset. Similar to Figure 3, (a) and (b) correspond to normalized respiration and CO_2_ recordings respectively and (c) shows their frequency spectra. Notice there is no low-frequency spectral mismatch in this clean data set.

### Neural Network

Obtaining the CO_2_ concentration from the respiration waveform is a 1D-to-1D (time series to time series) translation problem, which is modelled using a 1D fully convolutional encoder-decoder architecture. This modelling is analogous to prevalent image-to-image translation or semantic segmentation using 2D FCNs (Alotaibi, 2020; Long et al., 2015). However, all recent works in image-to-image translation problems involve adversarial training, which is notoriously hard especially with limited data. Thus, adversarial training is excluded in this paper.

Constructing a deep neural network often involves trial and error for tuning hidden layers. To find an optimum number of hidden layers in the network, several FCNs architectures are investigated, until overfitting was observed (test phase error increases with increasing network complexity). All codes are written in Python and use the PyTorch library, and would be publicly available on GitHub.

#### FCN Architecture

Input to the network was an array of size C x L, where the number of input channels, C = 1 and L is the length of recording. Although the respiration recordings were normalized using standard deviation, the resultant data range still varied between data sets. To bound the respiration amplitude within a fixed range, the respiration array was further normalized using the tanh operator before being passed on to the fully convolutional layers. We implemented four different FCN architectures, each having one (FCN-1L), two (FCN-2L), four (FCN-4L) and six (FCN-6L) convolution layers, respectively, between the input and output layers.

⍰ FCN-1L consisted of a single convolution operation with a kernel of length 7 and replicate padding of 3 on both sides (head and tail) of the input waveform. The kernel length is chosen to balance model complexity with accuracy.
⍰ FCN-2L encodes the tanh normalized respiration waveform by convolving it with a 4 × 7 kernel (4 kernels of length 7) with a stride of 2, which means the input is downsampled by a factor of 2. This is followed by ReLU nonlinearity (activation function) and finally a transposed convolution to decode the hidden layer into CO_2_. Both the convolution and transposed convolution are performed with a stride of 2, which replaces the need for a pooling layer to downsample the output of convolutional layers and an unpooling layer to upsample the output of transposed convolutional layers.
⍰ Similarly, FCN-4L consists of 2 convolution and 2 transposed convolutional layers, and FCN-6L architecture adds another 1 layer to both encoder and decoder sections. The network architecture of FCN-4L is shown in **Figure 5**.

**Figure 5.**
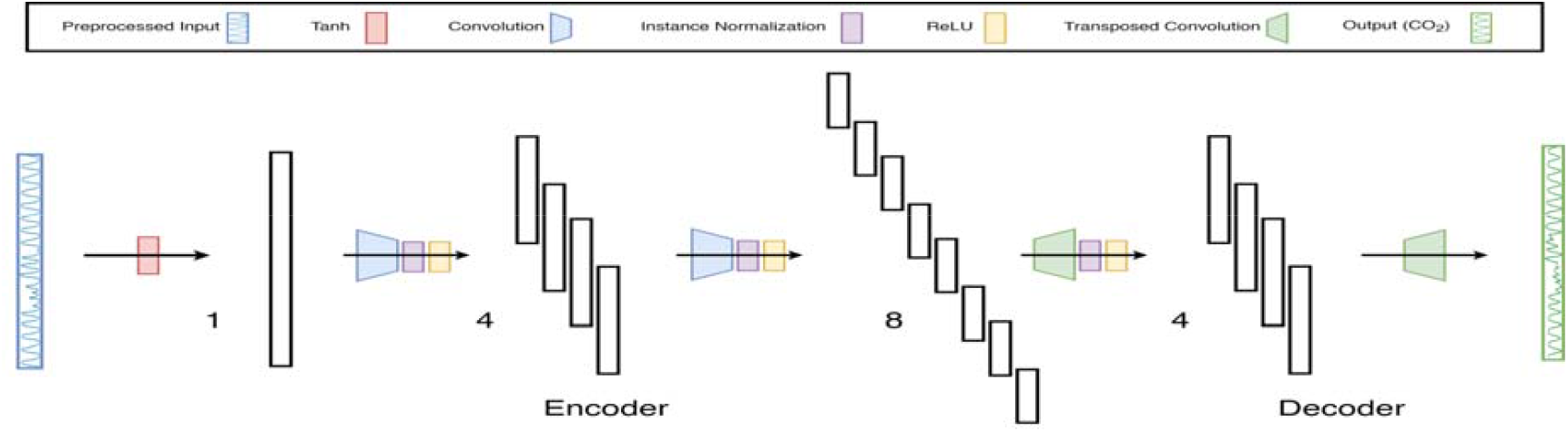
Neural-network architecture: 4-Layer Fully Convolutional Network. The architecture shown here is a type of encoder-decoder neural network consisting of fully convolutional layers, followed by instance normalization and ReLu non-linearity. The last layer doesn’t contain normalization and activation function as it is a regression problem. Moreover, the input is first normalized using tanh activation function to constrain the input data between -1 to 1.

#### Loss Function

We also experimented with two different loss functions. The first loss function is the mean squared error (MSE) computed between the measured and predicted CO_2_ waveforms, which is widely used in regression problems. However, as the regression was performed between the waveforms of pseudo-periodic nature, it was observed that the network learned to predict zero-crossings extremely well, but the extremities were left underfitted, lowering the scores of PETCO_2_ predictions. To rectify this problem, a second loss function, the weighted MSE, is introduced, with the weights set to the normalized amplitudes of the ground truth CO_2_ waveform for each timepoint. The weighting provides higher preference to the peaks, and hence we hypothesized that it would provide better results for PETCO_2_.

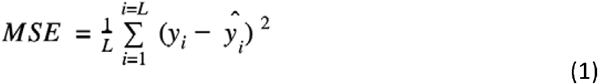

Where, ŷ_i_ and y_i_ are the predicted and ground truth CO_2_ respectively for the *i*^th^ time point, and L is the length of the recording. Networks trained with the weighted cost function are denoted by the postfix “-Wgt”.

#### Training

The 18 subjects were split into 5 subsets (splits), and the training was executed using the k-fold cross-validation strategy. It is typical to use either 10-fold or 5-fold cross-validation as it generally results in a model with low bias, modest variance and low-computational cost compared to leave-one-out cross-validation strategy (Rodriguez et al., 2010). In our dataset, as the number of subjects is relatively limited, we opted for k = 5, and each time one subset was left out from the training phase to be used in testing the accuracy of the network. Each subject can have multiple recordings, and the data was divided based on the subjects (and not recordings) to ensure that the training and testing data has no scans sharing a common subject. The divisions created by dotted lines in **Figure 2** correspond to the different splits. As visible in the figure, the splits contain data from 2, 5, 4, 4, 3 subjects, yielding total numbers of 30, 34, 27, 23, 22 recordings, respectively. Each split has a different number of total recordings, which enhances the generalizability of the results. We implemented two training strategies.

**Method 1: equal-length data segments**

In this method, we formatted the training data as an array of equal-sized data segments obtained by segmenting the input recordings. As the training is performed on a GPU, the computation can be parallelly performed in the tensor with multiple batches, reducing the training time. We used the chunk size of 90 seconds and a batch size of 256. The drawback of this method is the unavoidable error introduced due to edge effects during convolution, which is proportional to the number of chunks.

**Method 2: variable-length data segments**

In this method the input array length could be of variable sizes, as we are using a fully convolutional neural network. The drawback of using variable length input is that it prevents us from grouping the data in batches for parallel processing in the GPU. On the positive note, unlike the previous method, this training strategy precludes the segmenting-induced edge effects.

We implemented both methods. The training time was less than 20 seconds irrespective of the network type or training method. All the networks were trained using Adam optimizer for 15 epochs. Hyperparameters corresponding to the optimizer like learning rate and decay rate were fine tuned manually for each network. In total, we trained four FCNs, each using two loss functions, on the 5-fold split data. The training was performed on a 12GB GeForce GTX TITAN X GPU. All networks used less than 500MB GPU memory during the training phase.

#### Reference methods

To the best of our knowledge, there are no previous attempts to derive the CO_2_ waveform from respiratory traces using machine learning. To establish the performance of our approach against a possible alternative, we employed two reference methods. First, based on previous work by Chang et al., defining a PETCO_2_ as the convolution of RVT with RRF (and then normalized, negated and shifted temporally for maximum cross-correlation). This is referred to as the RVTRRF method, described by Eq. 2). RVT was estimated from respiration waveform as detailed in (Birn et al., 2008).

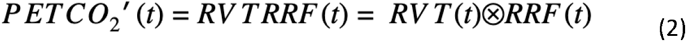

Where PETCO_2_’ is the estimated PETCO_2_. RRF is the respiratory response function. Similar to what was done previously (Chang and Glover, 2009), at the testing stage, we corrected the lag between RVTRRF (PETCO_2_′(t)) and PETCO_2_ using the maximum cross-correlation between the two signals, where the time shift was allowed to vary between -120 s and 120 s. Moreover, to maintain the scaling of PETCO_2_ as obtained from neural networks, we normalized and demeaned RVTRRF with the standard deviation and mean of PETCO_2_.

Second, defining a linear-regression (LR) model relating CO_2_ to respiratory volume (Eq. 3), and PETCO_2_’(t) is extracted from the CO_2_ time courses.

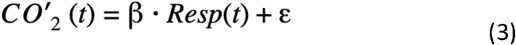

where CO_2_’ is the estimated CO_2_, є is the intercept, and β is the linear weighting factor derived from the “training data”, and the LR model could be understood as a single convolutional operation with a unit kernel size, making it similar to a machine learning linear regression problem. The training and testing partitioning are as described for the FCNs. MSE loss function was backpropagated similar to the FCNs.

#### Evaluation criteria

For the evaluation, the Pearson correlation coefficient (r), mean squared error (MSE), mean absolute error (MAE) and mean absolute percent error (MAPE) were calculated between (1) predicted CO_2_ and ground-truth CO_2_, (2) predicted PETCO2 and ground-truth PETCO_2_. Moreover, as the MAPE is sensitive to zero crossings, it was only calculated between the predicted PETCO_2_ and ground-truth PETCO_2_.

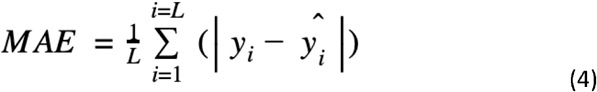

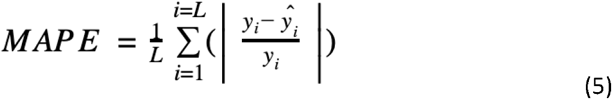

### Validation using functional MRI data

The final validation is inspired by a practical application of CO_2_ recordings, namely examining the relationship between PETCO_2_ and resting-state fMRI time series. For this we include data acquired from 2 healthy young subjects (male, age = 25 and 33 years). All data were acquired using a Siemens TIM Trio 3 T system and a 32-channel head coil. CO_2_ was acquired during these scans as described earlier.

⍰ Case 1: spin-echo EPI, TR = 323 ms, TE = 45 ms, flip angle = 90⍰, 2082 frames, voxel size = X: 3.48mm, Y: 3.48mm, Z: 6.25mm;
⍰ Case 2: gradient-echo EPI, TR = 323 ms, TE = 30 ms, 2230 frames, voxel size = : 3.48mm, Y: 3.48mm, Z: 6.25mm;
⍰ Case 3: simultaneous multi-slice gradient-echo EPI, TR = 323 ms, TE = 30 ms, flip angle = 40⍰, 2230 frames, voxel size = X: 3.48mm, Y: 3.48mm, Z: 6mm;

Preprocessing steps include: (1) filtering to 0.01 to 0.1 Hz band with AFNI (Cox, 1996); (2) spatial smoothing with a 5mm kernel (Jenkinson et al., 2012) (3) Discard the first 5 volumes in each scan to allow the brain to reach a steady state. All recorded and FCN-generated CO_2_ and PETCO_2_ time courses were downsampled to match the temporal resolution of the respective fMRI data.

## Results

Results for two representative data sets are shown in **Figure 6**. Method 1 (equal data length) adds no extra benefit to the training process and results in poor performance due to truncation effects in training data. Thus, all the results provided here are corresponding to method 2. The results are shown in **Figure 6** and summarized in Table 1. The best method, as determined by the lowest error terms (MSE, MAE, MAPE) and highest correlation (r) is identified by boldface. The predicted and ground truth PETCO_2_ signals show excellent visual agreement for FCN-4L-Wgt (**Figure 6B**). From Table 1, we can see that the CO_2_ error scores obtained from FCN-4L and FCN-4L-Wgt architecture are identical, with the scores corresponding to PETCO_2_ being slightly superior in the latter case. It should be noted that the Pearson correlation coefficient (r) remains unaffected by scaling and translation. Since the LR model involves only scaling and translation, the modeling step would not improve the baseline correlation between CO_2_ and respiration. Strangely, the RVTRRF model performs worse than the LR model (for PETCO_2_), suggesting that estimating PETCO_2_ from the peaks of the CO_2_ (and hence respiration) waveform may be more robust.

**Table 1.** Quantitative assessment of various network structures. RVTRRF = RVT convolved with RRF, LR = linear regression; FCN-XL = ‘X’ layered FCN used, -Wgt = with weighted MSE cost function. The parameters used in the assessment include: the correlation coefficient (r), the mean-squared error (MSE), the mean absolute error (MAE) and the mean-absolute percent error (MAPE). Each metric was calculated for every recording in the test set across all five splits. The mean and standard deviation (mean ± std) were calculated for all the metrics in each test split. Likewise, the average of (mean ± std) was taken across all the 5 splits and displayed in this table.

| Average scores across all five splits | RVTRRF | LR | FCN-1L | FCN-2L | FCN-4L | FCN-6L | FCN-4L-Wgt |
| --- | --- | --- | --- | --- | --- | --- | --- |
| r CO<sub>2</sub> | - | 0.901 ± 0.061 | 0.931 ± 0.055 | 0.922 ± 0.06 | **0.946 ± 0.054** | 0.944 ± 0.055 | **0.946 ± 0.056** |
| r PETCO<sub>2</sub> | 0.256 ± 0.132 | 0.311 ± 0.239 | 0.443 ± 0.261 | 0.45 ± 0.262 | 0.5 ± 0.266 | 0.461 ± 0.235 | **0.512 ± 0.269** |
| MSE CO<sub>2</sub> | - | 0.19 ± 0.103 | 0.138 ± 0.094 | 0.151 ± 0.101 | 0.108 ± 0.097 | 0.11 ± 0.097 | **0.106 ± 0.101** |
| MSE PETCO<sub>2</sub> | 0.032 ± 0.028 | 0.026 ± 0.021 | 0.02 ± 0.018 | 0.019 ± 0.017 | 0.018 ± 0.017 | 0.02 ± 0.018 | **0.017 ± 0.017** |
| MAE CO<sub>2</sub> | - | 0.337 ± 0.079 | 0.269 ± 0.077 | 0.276 ± 0.08 | 0.223 ± 0.076 | 0.227 ± 0.081 | **0.213 ± 0.08** |
| MAE PETCO<sub>2</sub> | 0.121 ± 0.055 | 0.112 ± 0.045 | 0.094 ± 0.04 | 0.093 ± 0.039 | 0.081 ± 0.035 | 0.085 ± 0.038 | **0.08 ± 0.036** |
| MAPE PETCO<sub>2</sub> | 0.125 ± 0.109 | 0.112 ± 0.084 | 0.094 ± 0.077 | 0.095 ± 0.078 | 0.085 ± 0.073 | 0.089 ± 0.074 | **0.084 ± 0.077** |

**Figure 6.**
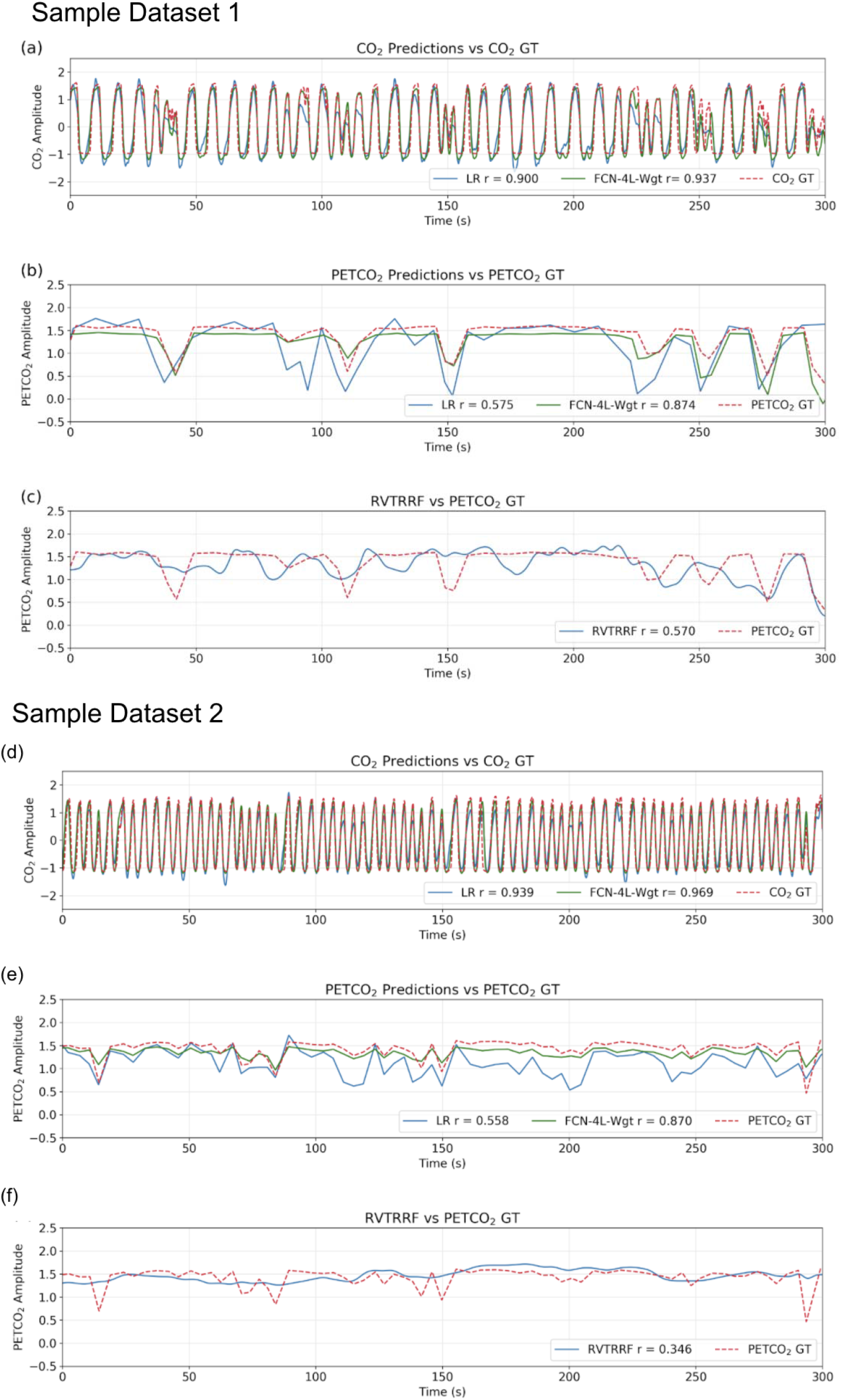
Qualitative comparison of resultant outputs. Two different sample predictions are shown from the test dataset, and for each of the example, comparisons are made between (a, d) the CO_2_ prediction and ground truth (GT), (b, e) the PETCO_2_ prediction from the reference linear regression model (LR), FCN-4L-Wgt model and the GT, and (c, f) PETCO_2_ estimated from RVTRRF and the PETCO_2_ GT.

Figure 7 shows the r score distribution across the entire test dataset for one of the five splits. The LR method is outperformed by all FCN methods for CO_2_ prediction. The difference between FCN-4L and its weighted counterpart is not noticable in the case of CO_2_ prediction, but overall, FCN-4L-Wgt achieved the highest r values, while FCN-6L achieved the lowest r-score variability. However, for PETCO_2_, the first and second quartile of FCN-4L-Wgt reach higher r values as compared to FCN-4L, making the superiority of weighted loss more evident. FCN-6L performs worse than all the other FCN networks in PETCO_2_ prediction. Note that the RVTRRF method is only able to reach a maximum r score of just below 0.5, substantially lower compared to all FCN networks. As previously mentioned, the r scores for RVTRRF correspond to maximum cross correlation with PETCO_2_, thus the scores are always positive. This step was not applied in other neural networks, hence for some of them the r scores between predicted and ground truth PETCO_2_ are negative.

**Figure 7.**
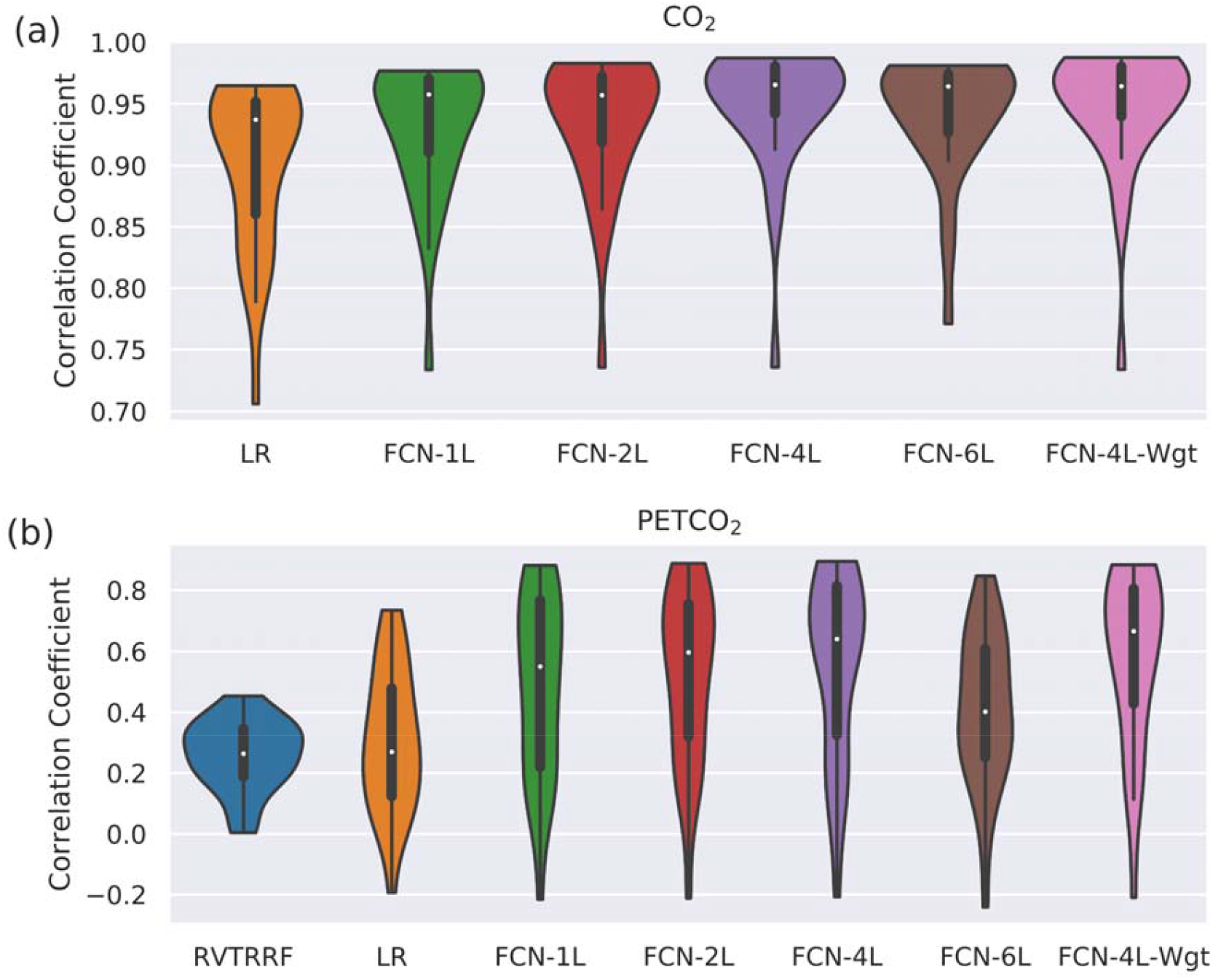
Performance of different methods: distribution of correlation coefficients (r) on test dataset. The r score is computed between (a) actual and predicted CO_2_, and (b) the actual and predicted PETCO_2_ obtained on the test dataset (for one of the five splits) is compared for different models used in the study and shown in the form of a bean plot. The median r for each method is shown as a white dot.

Figure 8 compares the correlation scores between training and testing phase for all the networks. From these plots, it can be inferred that FCN-6L likely overfits the training data, as reflected by a worse performance than that of the other networks (as reflected by a lower r). Since FCN-4L performs better than FCL-2L and doesn’t show huge differences between training and testing results, we can say four convolutional blocks are the optimum number for our given training data. Moreover, in our best model, MAPE score for PETCO_2_ is 0.142 (< 0.2), reflective of good prediction performance.

**Figure 8.**
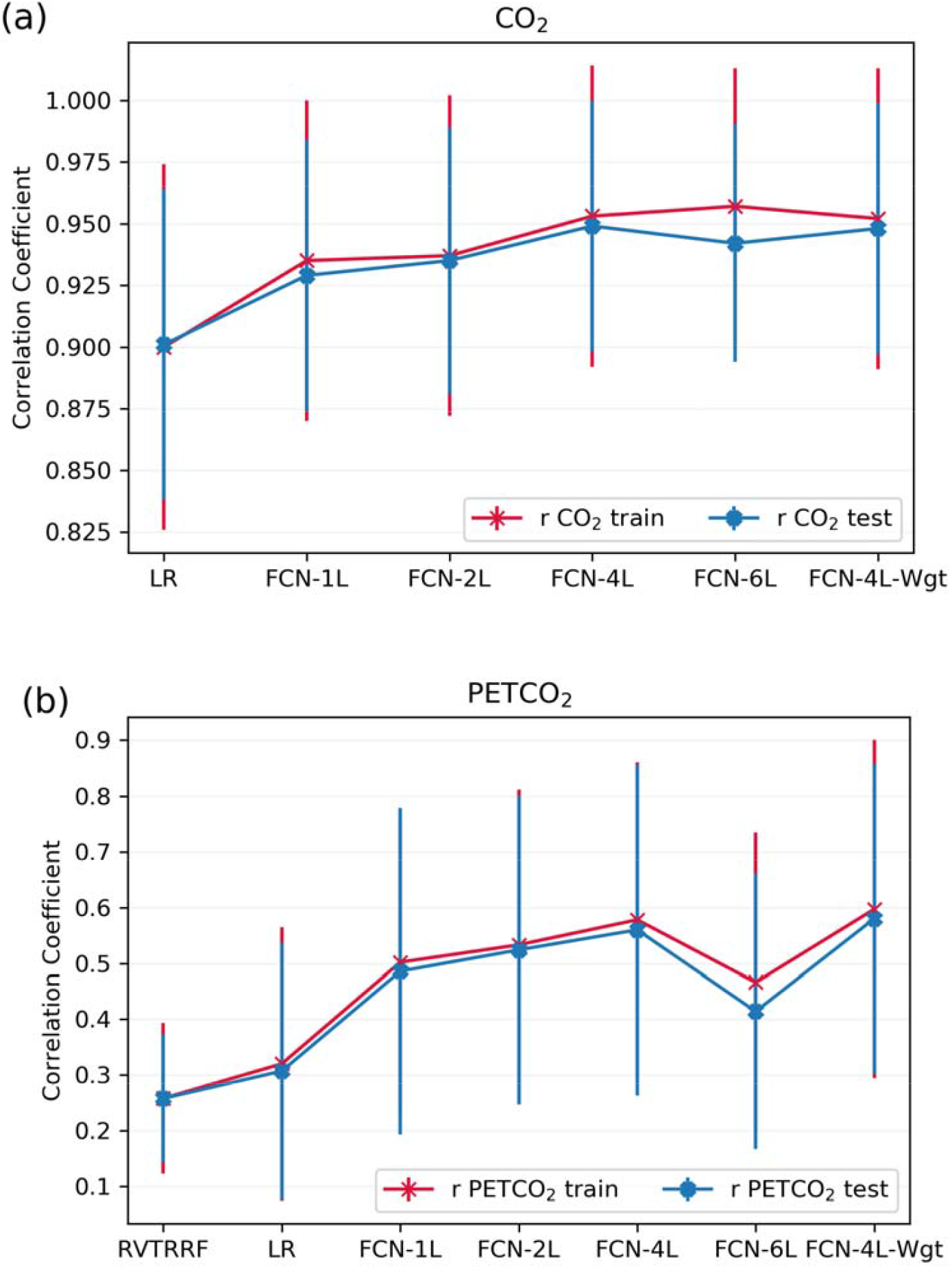
Comparison of model performance on train versus test datasets. The average Pearson correlation coefficient obtained across one of the splits for (a) CO_2_ and (b) PETCO_2_ between test and train dataset is shown in the top row. The error bars indicate the standard deviation.

Figure 9 compares the correlation scores across the five splits for all the networks. Perhaps counter-intuitively, the r-score ranking in the case of CO_2_ prediction does not match with that of PETCO_2_ prediction. In the case of CO_2_, the r scores of FCN-4L-Wgt closely resemble those of FCN-4L, but the former performed better for PETCO_2_ in all but one split. Though the best model varied depending on the split number and varies between CO_2_ and PETCO_2_ prediction, FCN-4L-Wgt consistently outperformed other models, exemplified in part by the highest correlation coefficients. The inter-split variability in r is the lowest for the reference methods (RVTRRF and LR) and highest for FCN methods, the various FCN methods themselves do not appear to exhibit different degrees of inter-split performance variability. Moreover, the performance rankings of the various methods are consistent across the splits and in line with the trends observed in **Figure 7**. Combining the results of **Figure 9** with the information in Figure 2, it can be seen that the poor performance in predicting CO_2_ across the 2nd split for all the models is due to one subject (subject 6). Split 3 performs the best in predicting CO_2_ across all the splits as the subjects in this split had higher correlation between CO_2_ and respiration compared to other splits. Still, the LR model performs worst in predicting PETCO_2_ in the 3rd split, reflecting that higher correlation between CO_2_ and respiration does not necessarily translate into higher correlation between peaks of CO_2_ and respiration. This point is further demonstrated by contrasting r scores of PETCO_2_ and CO_2_ for the LR approach in the remaining splits.

**Figure 9.**
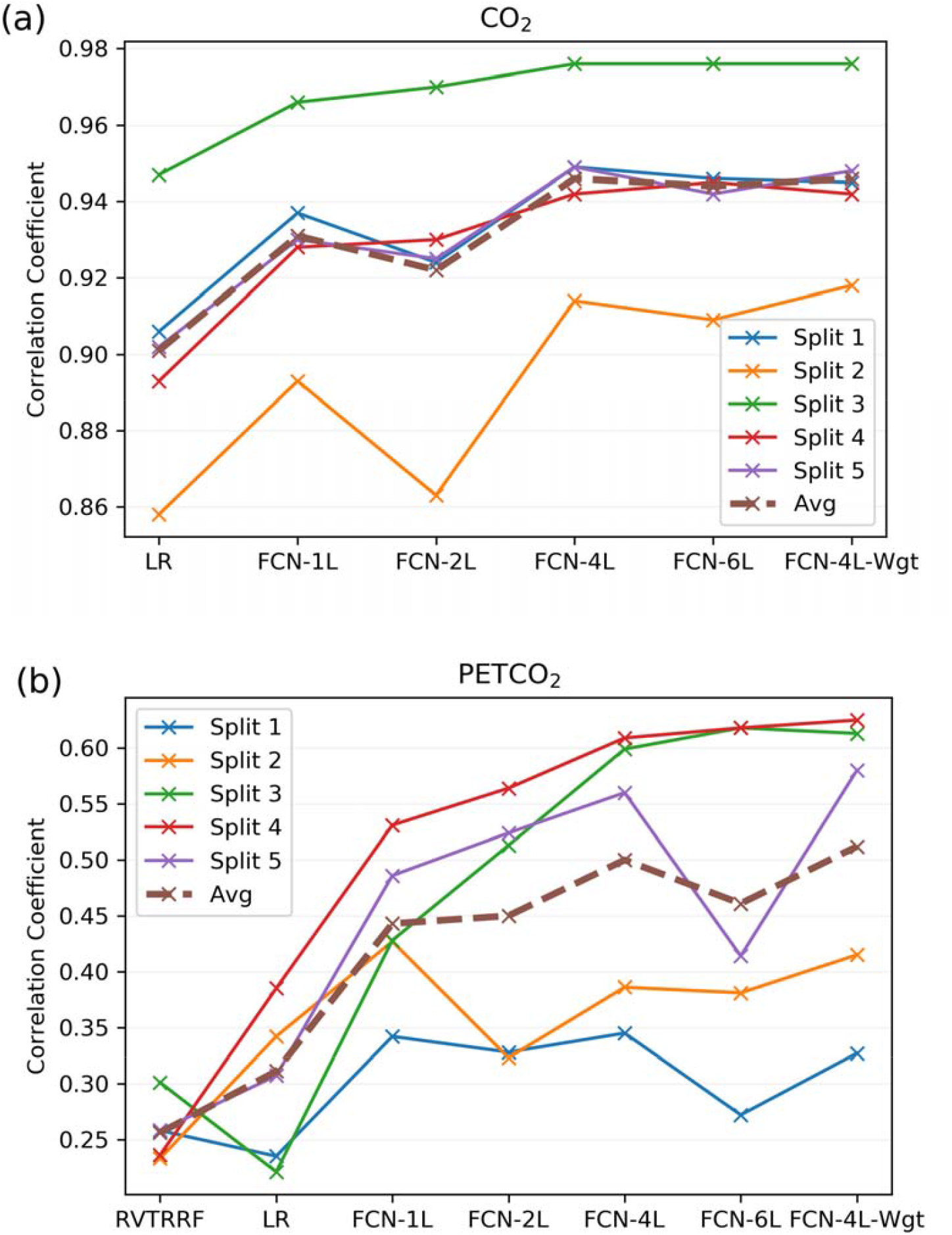
Model performance across the five splits. The correlation coefficients (r) obtained across the five splits and their average for all the models, for (a) CO_2_ and (b) PETCO_2_ prediction. The split number is the same as the splits shown in figure 2.

Figure 10 demonstrates the application of the FCN-4L-predicted PETCO_2_ time courses, which have established correlation with the resting-state fMRI signal. We show that the PETCO_2_-fMRI correlation maps for the ground-truth and predicted PETCO_2_ are highly similar in all scan sessions (Cases 1, 2 and 3) and subjects (Datasets 1 and 2). This preliminary demonstration suggests promise in using the model-predicted PETCO_2_.

**Figure 10.**
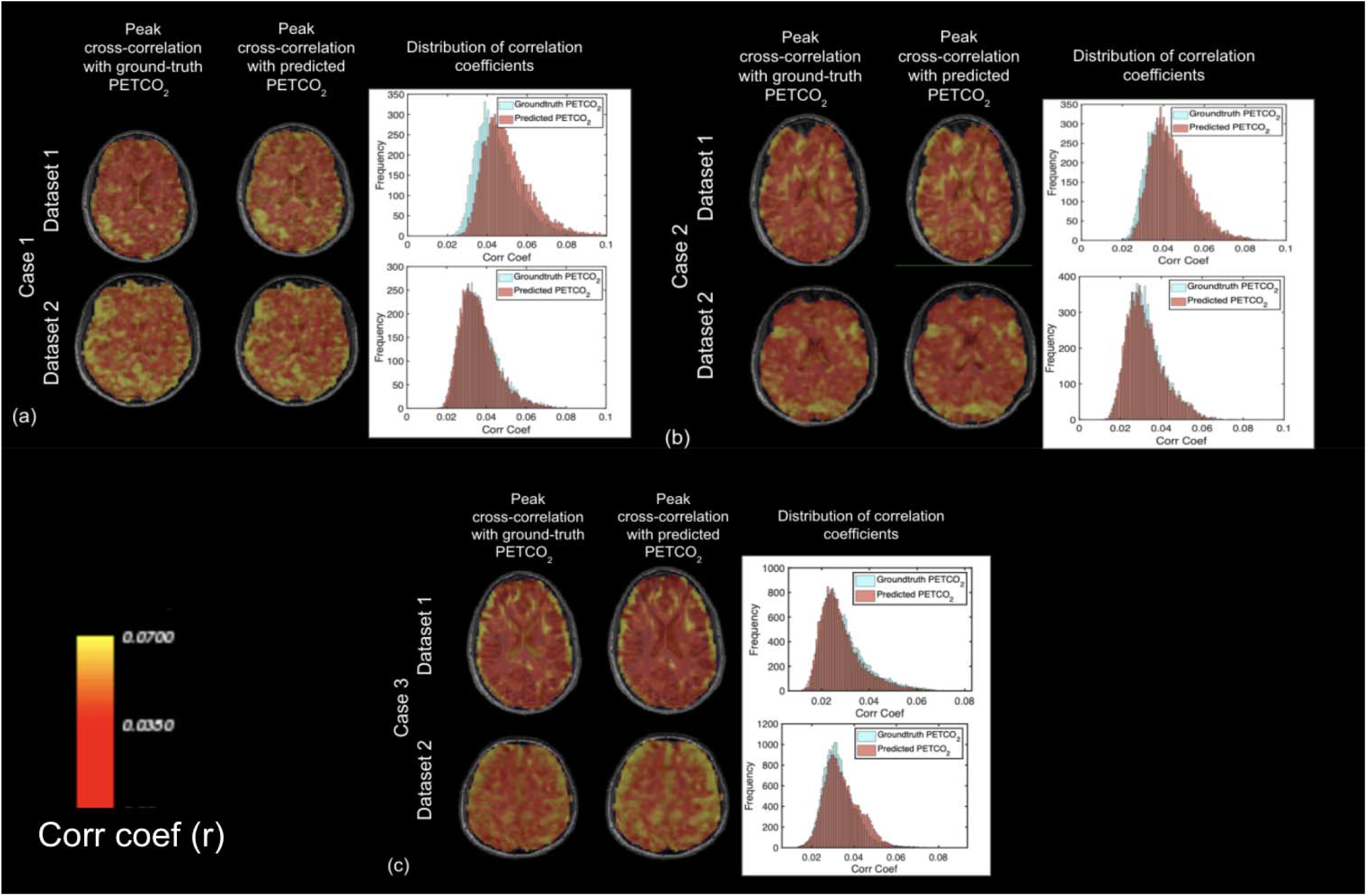
Comparison of ground-truth and predicted PETCO2 correlations. Data from 2 different subjects, imaged over multiple sessions (a, b, and c, respectively) are shown. In each case, the peak cross-correlation maps generated using the ground-truth and predicted PETCO_2_ time courses are compared side by side, with the corresponding correlation-coefficient histograms showing the comparability of the maps.

## Discussion

As a proof-of-concept study, we demonstrated that it is indeed feasible to use deep learning to predict CO_2_ from respiration. Furthermore, the performance of the FCN surpasses that of non-deep-learning methods. Note that the results only pertain to dynamic patterns in CO_2_, not to absolute CO_2_, which cannot be predicted from non-quantitative respiration patterns alone. Nonetheless, possible extensions and applications range from improving the feasibility of breath-holding based fMRI studies (Murphy et al., 2013) that lack CO_2_ recordings to the use of the CO2-O2 exchange ratio for vascular reactivity mapping (Chan et al., 2020, n.d.). In the following sections, we detail the machine-learning methodological considerations and possible future directions.

### Machine learning in physiological signal processing

The use of machine learning and deep learning models is prevalent in physiological signal data such as electromyogram (EMG), electroencephalogram (EEG), electrocardiogram (ECG), and electrooculogram (EOG) (Rim et al., 2020). It has been continuously observed that deep learning models perform better than other, classical machine learning models. Rim et al. conducted a review of 147 papers involving analysis using deep learning in EMG, ECG, EEG, EOG and their combinations (Rim et al., 2020). They concluded that in signal processing, most of the contributions were in the domain of classification task, feature-extraction task and data compression task, wherein CNN, RNN, CNN+RNN models were most commonly used. The complete pipeline was divided into 3 categories. The first category exploits machine learning models to extract features followed by DNN as a classifier to boost the accuracy of classification by obtaining useful features from raw data. The second type of pipeline involves deep learning model as a feature extractor and traditional machine learning model as a classifier to reduce hand-crafted labelling of the dataset. Finally, the third strategy uses an end-to-end deep learning pipeline to train raw data and receive the final output to build a robust DL model for the above-mentioned tasks. Due to the absence of a common study involving all 3 methods (Rim et al., 2020), we could not conclude on the best strategy. Our pipeline is positioned between the second and third categories, as we used an end-to-end DNN to estimate CO_2_ as an intermediate step, followed by a post-processing step to obtain the final PETCO_2_ waveform.

### Utility and Current Status of using RVT for generating PETCO_2_

As RVTRRF is correlated with PETCO_2_, there is a potential of training a convolutional neural network between RVT and PETCO_2_, which might perform better than a single convolution operation using RRF. This approach appeals to find a neural network architecture which could replace the need of RRF. We experimented with different types of neural networks trained to predict PETCO_2_ from RVT. However, during our analysis, we found that none of the networks could adequately generate PETCO_2_ from RVT. We assert that, even if there were sufficient dataset for the training, this approach is expected to always perform worse compared to training the neural network to associate respiration and CO_2_. Our rationale is the fact that the latter exploits the evident breathing pattern between respiration patterns and CO_2_ and performs well even with limited recording lengths. Conversely, in the former approach, the temporal resolution of RVT is fundamentally constrained to the observed breath durations, and the peak detection algorithm can often miss deep breaths (Power et al., 2020). Therefore, using RVT to predict PETCO_2_ may not be robust compared to directly using respiration belt recordings.

As a potential alternative metric of respiratory variability, the windowed respiratory variance (RV), computed as the standard deviation of the respiratory signal over sliding windows of 6 s (Chang et al., 2009), is more robust than RVT against noise. It is intuitively appreciated, as RV is taken over all respiratory cycles over its sliding window rather than being based on the depth-to-duration ratio (as in the case of RVT). Removing the influence of breath-cycle duration term may render RV less physiologically related to CO_2_ than RVT, and we did not investigate the use of RV in this proof-of-concept study because the analytical RRF we use is defined from RVT (Birn et al., 2008). Another potential influence on the performance of CO_2_ prediction may be the presence of hardware/software bandpass filters on the raw recordings. The Biopac system provided software filters to exclude MRI noise (periodicity < 100 ms) while preserving higher frequencies, and it is conceivable that in cases where such frequencies are removed from the raw respiratory traces, the ability to predict CO_2_ fluctuations may be disadvantaged.

### Other Deep Learning Architectures

We are aware of alternative, recently developed network architectures that may suit our problem. For instance, unpaired and paired image-to-image translation tasks have been accomplished by generative adversarial networks (GANs) such as Pix2Pix (Isola et al., 2017) and CycleGAN (Zhu et al., 2017). The task of transforming the respiration belt data to the CO_2_ waveform is analogous to paired image-to-image translation. A simple GAN consists of two sub-models, a generator to obtain synthetic samples, and a discriminator that predicts the truth value of the provided sample; trained together in a zero-sum game, such that discriminator’s guess is no better than random and the generator wins at the end of training. The discriminator network in GANs is similar to the explicit loss function used in traditional deep learning models. In our case, adversarial training would mean that instead of using a MSE or weighted MSE loss function to determine the best CO_2_ prediction, another network would distinguish between them. Given that our use case is much simpler, this approach might not add any extra value at the cost of computation and potential overfitting.

Another category of deep learning models is the recurrent neural networks (RNNs), such as the long-short term memory (LSTM) (Greff et al., 2017) and gated recurrent unit (GRU) (Zhao et al., 2016) networks, which are widely used for time series analysis and signal processing. At first glance, RNNs seemed to be suited to our use case, but unfortunately, we could not obtain good results using RNNs. Specifically, as an alternative to the FCN, we had implemented a convolutional LSTM, supplied with an input of 5 seconds respiration data so that we would capture at least the waveform from one breath. The initial 5-second respiration-signal segment was fed into the LSTM block which would predict the corresponding segment of CO_2_ and the hidden state. These outputs along with the next 5-second segment of respiration data were used as the inputs for the next iteration, with the intention that irregularities in breathing would be stored in the network’s memory and would help in prediction. Unfortunately, because of the short input-lengths, coupled with the limited durations of respiration recordings, the concatenated output lacked the smooth transition between consecutive chunks (exhibiting edge effects in each 5-second block, similar to observed in training method 1), which are required for accurately predicting a slow-varying signal like PETCO_2_. Thus, we concluded that time-series to time-series translation using RNNs is not feasible unless much longer respiratory and CO_2_ recordings were available.

## Limitations

The sources of error, which might be biasing our results, can come from the errors in our dataset. In our opinion, the quality assurance section takes care of it by rejecting such recordings. Another possibility of biases in our results is the split of test and train data, but 5-fold cross-validation reduces such bias. One last source of error is the peak detection algorithm. Currently, the scores corresponding to PETCO_2_ comparison include errors introduced by the peak detection algorithm applied on CO_2_ data. A perfect peak detection algorithm might provide us with slightly better scores.

In this work, we demonstrated the estimation of dynamic PCO_2_ and PETCO_2_ time-series estimation, but our method does not attach quantitative values to either metric (e.g. in units of mmHg). This is mainly because we expect quantitative end-tidal CO_2_ values obtained from respiratory-belt recordings to be inaccurate. This is likely due to the fact that the extent to which spontaneous PETCO_2_ reflects arterial CO_2_ depends on alveolar tension, which in turn depends on minute ventilation (Rawat et al., 2021). Minute ventilation, in turn, depends on respiratory rate and tidal volume (Quanjer et al., 1993; Rawat et al., 2021; Stocks and Quanjer, 1995), which do not instantaneously translate into PETCO_2_. Indeed, in our data, we did not find a one-to-one relationship between point-by-point PETCO_2_ and respiratory depth variations. Thus, our breath-by-breath CO_2_ time series reflects patterns of change but not quantitative values of PETCO_2_, and the utility of our neural network remains focused on the need for dynamic variations in PCO_2_ and PETCO_2_.

## Conclusion

This study demonstrates the possibility of reconstructing PETCO_2_ data from respiratory recordings, thus, enabling broader incorporation of PETCO_2_ in rs-fMRI analysis. Borrowing the analogy from image-to-image translation using 2D FCNs, we introduced 1D FCNs for 1D signal to signal translation. The trained neural network outperformed the linear regression and RVTRRF models. The study also contrasts the depth of the FCNs. As we hypothesized, the network trained using weighted MSE performs better for a FCN architecture compared to MSE loss function. The results across different deep neural network architectures serve as a literature for further research in signal processing and for the deep learning community.

## Acknowledgment

The authors acknowledge funding support from the Canadian Institutes of Health Research (CIHR) and the Natural Sciences and Engineering Research Council of Canada (NSERC). We also thank Dr. Catie Chang (Vanderbilt) for helpful comments.

## References

Alotaibi, A., 2020. Deep Generative Adversarial Networks for Image-to-Image Translation: A Review. Symmetry 12, 1705. https://doi.org/10.3390/sym12101705

Bayrak, R.G., Salas, J.A., Huo, Y., Chang, C., 2020. A Deep Pattern Recognition Approach for Inferring Respiratory Volume Fluctuations from fMRI Data, in: Medical Image Computing and Computer Assisted Intervention – MICCAI 2020. Springer International Publishing, pp. 428–436. https://doi.org/10.1007/978-3-030-59728-3_42

Birn, R.M., Diamond, J.B., Smith, M.A., Bandettini, P.A., 2006. Separating respiratory-variation-related fluctuations from neuronal-activity-related fluctuations in fMRI. Neuroimage 31, 1536–1548.

Birn, R.M., Smith, M.A., Jones, T.B., Bandettini, P.A., 2008. The respiration response function: the temporal dynamics of fMRI signal fluctuations related to changes in respiration. Neuroimage 40, 644–654. https://doi.org/10.1016/j.neuroimage.2007.11.059

Blockley, N.P., Harkin, J.W., Bulte, D.P., 2017. Rapid cerebrovascular reactivity mapping: Enabling vascular reactivity information to be routinely acquired. Neuroimage. https://doi.org/10.1016/j.neuroimage.2017.07.048

Bright, M.G., Whittaker, J.R., Driver, I.D., Murphy, K., 2020. Vascular physiology drives functional brain networks. Neuroimage 116907. https://doi.org/10.1016/j.neuroimage.2020.116907

Champagne, A.A., Bhogal, A.A., Coverdale, N.S., Mark, C.I., Cook, D.J., 2019. A novel perspective to calibrate temporal delays in cerebrovascular reactivity using hypercapnic and hyperoxic respiratory challenges. NeuroImage. https://doi.org/10.1016/j.neuroimage.2017.11.044

Chang, C., Cunningham, J.P., Glover, G.H., 2009. Influence of heart rate on the BOLD signal: the cardiac response function. Neuroimage 44, 857–869.

Chang, C., Glover, G.H., 2009. Relationship between respiration, end-tidal CO2, and BOLD signals in resting-state fMRI. Neuroimage 47, 1381–1393. https://doi.org/10.1016/j.neuroimage.2009.04.048

Chan, S.-T., Evans, K.C., Song, T.-Y., Selb, J., van der Kouwe, A., Rosen, B.R., Zheng, Y.-P., Ahn, A., Kwong, K.K., 2020. Cerebrovascular reactivity assessment with O2-CO2 exchange ratio under brief breath hold challenge. PLoS One 15, e0225915. https://doi.org/10.1371/journal.pone.0225915

Chan, S.T., Ordway, C., Calvanio, R.J., Buonanno, F.S., Rosen, B.R., Kwong, K.K., n.d. Cerebrovascular responses to O2-CO2 exchange ratio under brief breath-hold challenge in patients with chronic mild traumatic brain injury. https://doi.org/10.1101/2021.04.22.441010

Chen, J.J., 2018. Cerebrovascular-Reactivity Mapping Using MRI: Considerations for Alzheimer’s Disease. Front. Aging Neurosci.

Chen, J.J., Gauthier, C.J., 2021. The Role of Cerebrovascular-Reactivity Mapping in Functional MRI: Calibrated fMRI and Resting-State fMRI. Front. Physiol. 12, 657362. https://doi.org/10.3389/fphys.2021.657362

Cox, R.W., 1996. AFNI: software for analysis and visualization of functional magnetic resonance neuroimages. Comput. Biomed. Res. 29, 162–173. https://doi.org/10.1006/cbmr.1996.0014

Golestani, A.M., Chang, C., Kwinta, J.B., Khatamian, Y.B., Chen, J.J., 2015. Mapping the end-tidal CO2 response function in the resting-state BOLD fMRI signal: Spatial specificity, test– retest reliability and effect of fMRI sampling rate. Neuroimage 104, 266 – 277. https://doi.org/10.1016/j.neuroimage.2014.10.031

Golestani, A.M., Chen, J.J., 2020. Controlling for the effect of arterial-CO2 fluctuations in resting-state fMRI: Comparing end-tidal CO2 clamping and retroactive CO2 correction. Neuroimage 116874. https://doi.org/10.1016/j.neuroimage.2020.116874

Greff, K., Srivastava, R.K., Koutnik, J., Steunebrink, B.R., Schmidhuber, J., 2017. LSTM: A Search Space Odyssey. IEEE Trans Neural Netw Learn Syst 28, 2222–2232. https://doi.org/10.1109/TNNLS.2016.2582924

Hülsmann, W.C., Dubelaar, M.L., 1988. Aspects of fatty acid metabolism in vascular endothelial cells. Biochimie 70, 681–686. https://doi.org/10.1016/0300-9084(88)90253-2

Iadecola, C., 2017. The Neurovascular Unit Coming of Age: A Journey through Neurovascular Coupling in Health and Disease. Neuron 96, 17–42. https://doi.org/10.1016/j.neuron.2017.07.030

Isola, P., Zhu, J.-Y., Zhou, T., Efros, A.A., 2017. Image-to-Image Translation with Conditional Adversarial Networks. 2017 IEEE Conference on Computer Vision and Pattern Recognition (CVPR). https://doi.org/10.1109/cvpr.2017.632

Jenkinson, M., Beckmann, C.F., Behrens, T.E.J., Woolrich, M.W., Smith, S.M., 2012. FSL. Neuroimage 62, 782–790. https://doi.org/10.1016/j.neuroimage.2011.09.015

Kaji, S., Kida, S., 2019. Overview of image-to-image translation by use of deep neural networks: denoising, super-resolution, modality conversion, and reconstruction in medical imaging. Radiol. Phys. Technol. 12, 235–248. https://doi.org/10.1007/s12194-019-00520-y

Kiranyaz, S., Avci, O., Abdeljaber, O., Ince, T., Gabbouj, M., Inman, D.J., 2021. 1D convolutional neural networks and applications: A survey. Mechanical Systems and Signal Processing. https://doi.org/10.1016/j.ymssp.2020.107398

Komori, M., Takada, K., Tomizawa, Y., Nishiyama, K., Kawamata, M., Ozaki, M., 2007. Permissive range of hypercapnia for improved peripheral microcirculation and cardiac output in rabbits. Crit. Care Med. 35, 2171–2175. https://doi.org/10.1097/01.ccm.0000281445.77223.31

Long, J., Shelhamer, E., Darrell, T., 2015. Fully convolutional networks for semantic segmentation. 2015 IEEE Conference on Computer Vision and Pattern Recognition (CVPR). https://doi.org/10.1109/cvpr.2015.7298965

Murphy, K., Birn, R.M., Bandettini, P.A., 2013. Resting-state fMRI confounds and cleanup. Neuroimage 80, 349–359. https://doi.org/10.1016/j.neuroimage.2013.04.001

Najarian, T., Marrache, A.M., Dumont, I., Hardy, P., Beauchamp, M.H., Hou, X., Peri, K., Gobeil, F., Jr, Varma, D.R., Chemtob, S., 2000. Prolonged hypercapnia-evoked cerebral hyperemia via K(+) channel- and prostaglandin E(2)-dependent endothelial nitric oxide synthase induction. Circ. Res. 87, 1149–1156. https://doi.org/10.1161/01.res.87.12.1149

Nikolaou, F., Orphanidou, C., Papakyriakou, P., Murphy, K., Wise, R.G., Mitsis, G.D., 2016. Spontaneous physiological variability modulates dynamic functional connectivity in resting-state functional magnetic resonance imaging. Philos. Trans. A Math. Phys. Eng. Sci. 374. https://doi.org/10.1098/rsta.2015.0183

Peebles, K., Celi, L., McGrattan, K., Murrell, C., Thomas, K., Ainslie, P.N., 2007. Human cerebrovascular and ventilatory CO2 reactivity to end-tidal, arterial and internal jugular vein PCO2. J. Physiol. 584, 347–357. https://doi.org/10.1113/jphysiol.2007.137075

Peebles, K.C., Richards, A.M., Celi, L., McGrattan, K., Murrell, C.J., Ainslie, P.N., 2008. Human cerebral arteriovenous vasoactive exchange during alterations in arterial blood gases. J. Appl. Physiol. 105, 1060–1068. https://doi.org/10.1152/japplphysiol.90613.2008

Pelligrino, D.A., Santizo, R.A., Wang, Q., 1999. Miconazole represses CO(2)-induced pial arteriolar dilation only under selected circumstances. Am. J. Physiol. 277, H1484–90. https://doi.org/10.1152/ajpheart.1999.277.4.H1484

Pinto, J., Bright, M.G., Bulte, D.P., Figueiredo, P., 2020. Cerebrovascular Reactivity Mapping Without Gas Challenges: A Methodological Guide. Front. Physiol. 11, 608475. https://doi.org/10.3389/fphys.2020.608475

Power, J.D., Lynch, C.J., Dubin, M.J., Silver, B.M., Martin, A., Jones, R.M., 2020. Characteristics of respiratory measures in young adults scanned at rest, including systematic changes and “missed” deep breaths. Neuroimage 204, 116234. https://doi.org/10.1016/j.neuroimage.2019.116234

Quanjer, P.H., Tammeling, G.J., Cotes, J.E., Pedersen, O.F., Peslin, R., Yernault, J.C., 1993. Lung volumes and forced ventilatory flows. Eur. Respir. J. 6 Suppl 16, 5–40. https://doi.org/10.1183/09041950.005s1693

Rawat, D., Modi, P., Sharma, S., 2021. Hypercapnea, in: StatPearls. StatPearls Publishing, Treasure Island (FL).

Rawat, W., Wang, Z., 2017. Deep Convolutional Neural Networks for Image Classification: A Comprehensive Review. Neural Comput. 29, 2352–2449. https://doi.org/10.1162/NECO_a_00990

Rim, B., Sung, N.-J., Min, S., Hong, M., 2020. Deep Learning in Physiological Signal Data: A Survey. Sensors. https://doi.org/10.3390/s20040969

Rodriguez, J.D., Perez, A., Lozano, J.A., 2010. Sensitivity Analysis of k-Fold Cross Validation in Prediction Error Estimation. IEEE Transactions on Pattern Analysis and Machine Intelligence. https://doi.org/10.1109/tpami.2009.187

Salas, J.A., Bayrak, R.G., Huo, Y., Chang, C., 2020. Reconstruction of respiratory variation signals from fMRI data. NeuroImage. https://doi.org/10.1016/j.neuroimage.2020.117459

Stocks, J., Quanjer, P.H., 1995. Reference values for residual volume, functional residual capacity and total lung capacity. ATS Workshop on Lung Volume Measurements. Official Statement of The European Respiratory Society. Eur. Respir. J. 8, 492–506. https://doi.org/10.1183/09031936.95.08030492

Wise, R.G., Ide, K., Poulin, M.J., Tracey, I., 2004. Resting fluctuations in arterial carbon dioxide induce significant low frequency variations in BOLD signal. Neuroimage 21, 1652–1664.

Zhao, Y., Li, J., Xu, S., Xu, B., 2016. Investigating gated recurrent neural networks for acoustic modeling. 2016 10th International Symposium on Chinese Spoken Language Processing (ISCSLP). https://doi.org/10.1109/iscslp.2016.7918370

Zhu, G., Jiang, B., Tong, L., Xie, Y., Zaharchuk, G., Wintermark, M., 2019. Applications of Deep Learning to Neuro-Imaging Techniques. Front. Neurol. 10. https://doi.org/10.3389/fneur.2019.00869

Zhu, J.-Y., Park, T., Isola, P., Efros, A.A., 2017. Unpaired Image-to-Image Translation Using Cycle-Consistent Adversarial Networks. 2017 IEEE International Conference on Computer Vision (ICCV). https://doi.org/10.1109/iccv.2017.244

